# Bio-guided isolation of a new sesquiterpene from *Artemisia cina* with anthelmintic activity against *Haemonchus contortus* L_3_ infective larvae

**DOI:** 10.1101/2024.02.26.582230

**Authors:** Luis David Arango-De-la Pava, Manasés González-Cortázar, Alejandro Zamilpa, Jorge Alfredo Cuéllar-Ordaz, Héctor Alejandro de la Cruz-Cruz, Rosa Isabel Higuera-Piedrahita, Raquel López-Arellano

## Abstract

*Haemonchus contortus* is a blood-feeding gastrointestinal parasite that impacts grazing sheep, causing economic losses in animal production and presenting anthelmintic resistance, requiring alternative antiparasitic treatments, including the exploration of plant-based anthelmintics. *Artemisia cina* (Asteraceae) is a plant whose *n*-hexane (*n*-HE) and ethyl acetate extract (EAE) exhibits anthelmintic activity against *H. contortus*, being the *n*-HE more active. With the aim of discovering additional bioactive metabolites, a chemical analysis was performed on EAE, which presented a LC_90_ of 3.30 mg/mL and allowed the isolation of 11-[(1*R*,5*S*,7*R*,8*R*,10*S*,)-1,8-dihydroxy-5,10-dimethyl-4-oxodecahydroazulen-7-yl] acrylic acid, a new sesquiterpene that was identified through one and two-dimensional NMR. The compound was named cinic acid and displayed a CL_50_ of 0.13 (0.11 -0.14) mg/mL and CL_90_ of 0.40 (0.37 - 0.44) mg/mL, which, compared with EAE larvicidal activity, was 256-fold more active at LC_50_ and 15.71-fold at LC_90_. In this study, a new sesquiterpene with anthelmintic effects against *H. contortus* L_3_ infective larvae was isolated from the EAE of *Artemisia cina*.

**Author summary:** *Haemonchus contortus* is a hematophagous gastrointestinal parasite that affects grazing sheep. Due to its feeding habits, it induces anemia, poor digestion, diarrhea, and weight loss in animals, potentially leading to death in young individuals and causing economic losses in animal production. Moreover, it demonstrates resistance to drugs, making it imperative to explore alternative antiparasitic treatments against *H. contortus*, such as the discovery and development of plant-based anthelmintics. In this work, we explore the ethyl acetate extract (EAE) of *Artemisia cina* in the search of bioactive compounds. A new sesquiterpene was separated through a bio-guided isolation monitoring the larvicidal effect against *H. Contortus* L_3_ infective larvae and was named cinic acid. These findings suggest that the EAE could be promising candidate for the development of a plant-based pharmaceutical preparation with notable anthelmintic activity against *H. contortus*.

## Introduction

*Haemonchus contortus* is a highly pathogenic nematode that feeds on the blood of small ruminants and is a significant cause of economic losses worldwide. It possesses a particularly significant threat in tropical, subtropical, and warm temperate regions where warm and moist conditions favor the free-living stages of the parasite [1]. Females of *H. contortus* can produce up to 5,000 eggs per day, which are then excreted from the host animal through feces. After hatching, the larvae undergo several chitin molts, ultimately reaching an infective larval stage known as L_3_. This larva is ingested by ruminants as they consume grass. Upon reaching the abomasum, the L_4_ larva initiates its blood-feeding role, potentially causing a range of issues such as malnutrition, low feed conversion, anemia, loss of appetite, low fertility rates, and even death in both young and older animals. Chronic inflammation, weight loss, and continuous diarrhea may contribute to the deterioration of the animal’s health and ultimately lead to its demise [2].

The control of *H. contortus* in small ruminants has been achieved using chemical anthelmintics [3]. However, their inadequate and irresponsible use has facilitated the emergence of parasites with resistance to anthelmintics in different countries, including Mexico, where small ruminant grazing is a significant economic activity [4, 5]. Therefore, it is imperative to explore and propose alternative control strategies for this parasite. Among the various options, the use of plant extracts containing chemical compounds with anthelmintic activity holds promise.

The genus *Artemisia* comprises approximately 500 species, distributed worldwide. *Artemisia* species are characterized as small herbs or shrubs with a distinctive bitter taste and a pungent aroma attributed to terpenoids, primarily monoterpenes in the essential oil, and sesquiterpene lactones[6]. They also, comprise terpenoids, flavonoids, coumarins, caffeoylquinic acids, sterols and acetylenes[7]. *Artemisia cina*, also known as santonica or Levant wormseed, has been traditionally used as a vermifuge to expel intestinal worms. The efficacy of *A. cina* against *H. contortus* has been demonstrated both *in vitro* and *in vivo. In vitro*, the *n*-hexane (*n*-HE) extract of *A. cina* exhibited the highest larvicidal activity against transitional larvae L_3_-L_4_ of *H. contortus* compared to methanol and ethyl acetate extracts (EAE), achieving percentages of 75% and 82.6% at concentrations of 1 mg/mL and 2 mg/mL, respectively[4]. In an *in vivo* study conducted on naturally infected periparturient goats, the administration of an *n*-HE derived from *A. cina* resulted in a notable reduction in the fecal egg count of *H. contortus* and *Teladorsagia circumcincta*. This extract was found to contain two previously unidentified compounds for *A. cina*, namely isoguaiacin and norisoguaiacin [8].

According to previous findings, the EAE presents anthelmintic activity against *H. contortus* L_3_ infective larvae [8], however the bioactives compounds remain unidentified. Therefore, the objective of this study was to isolate and identify a compound with anthelmintic activity against *H. contortus* L_3_ infective larvae from the EAE of *A. cina* through bio-guided separation. This is the first time that the 11-[(1*R*,5*S*,7*R*,8*R*,10*S*,)-1,8-dihydroxy-5,10-dimethyl-4-oxodecahydroazulen-7-yl] acrylic acid (cinic acid) and its anthelmintic activity against L_3_ *H. contortus* infective larvae is reported.

## Materials and methods

### Plant Material

The fresh pre-flowering leaves and stems of *A. cina* O. Berg ex Poljakov (Asteraceae) (10 kg) were bought at Hunab® laboratory. A voucher specimen was authenticated by Dr. Alejandro Torres-Montúfar and was deposited at the herbarium of Facultad de Estudios Superiores Cuautitlán (FES-C) UNAM, México under voucher no 11967. The plant was grown at 80% humidity, 24 °C temperature and soil with pH = 6.3.

### *Artemisia cina* extract

The extracts were obtained by maceration. Dry *A. cina* leaves and stems (1 kg) were ground and placed in 1 L erlenmeyers. Extraction using leaves and stems was performed using *n*-hexane, ethyl acetate and methanol maintained for 72 h at room temperature (23–25 °C). The extraction was performed using fresh vegetal material for each solvent, avoiding exhaustive extraction method used by Higuera-Piedrahita et al. 2021 [4]. Extracts were filtered using a Whatman No. 4 paper and the solvent were removed by low-pressure distillation using a rotary evaporator (DLAB RE-100 Pro) at 40 °C and 100 rpm. The extract was finally lyophilized and kept at 4 °C for phytochemical and biological assays.

### *In vitro* assays with *Haeomonchus contortus* L_3_

The lethal effect of the three *A. cina* extracts and fractions on L_3_ *H. contortus* infective larvae was determined using 96-well microplates for 24 hours at 24 °C. Two control groups were used: (a) Distilled water and (b) ivermectin (5 mg/mL, Sigma-Aldrich). *A. cina* extracts were tested at five different concentrations (8, 4, 2, 1, and 0.5mg/mL). For the *A. cina* extract × *H. contortus* infective larvae confrontations, approximately 100 L_3_ larvae in 100 μL of aqueous suspension were used per well (n = 4), with three replicates under the same conditions. The lethal effect was evaluated at 24 hours post-exposure, and lethality percentages were obtained. The *H. contortus* strain was obtained from Facultad de Estudios Superiores Cuautitlán.

### Bio-guided separation

24 g of the (ethyl acetate extract) EAE were utilized for column 1 and separated using open column chromatography. Normal silica gel 60 (Merck®, 0.015-0.040 mm) served as stationary phase and *n*-hexane-ethyl acetate was employed as solvent gradient system. 61 samples were obtained and grouped in three fractions according to their chemical similarity, that was monitored using thin-layer chromatography. Samples were concentrated using a rotary evaporator. The resulting fractions were named C1F1 (4.242 g), C1F2 (11.187 g) and C1F3 (5.691 g).

C1F2 achieved the highest yield percentage with larvicidal activity, so it was used for column 2. Column 2 was performed using the same chromatographic conditions as Column 1. 34 samples were grouped into nine fractions: C2F1 (0.093 g), C2F2 (0.088 g), C2F3 (0.090 g), C2F4 (0.137 g), C2F5 (1.727 g), C2F6 (2.656 g), C2F7 (2.780 g), C2F8 (0.769 g), and C2F9 (1.142 g). C2F1 to C-F4 were not evaluated due to low yield percentages Instead, C2F5 to C2F7 were analyzed at C1F2 CL_50_ and CL_90_. C2F7 displayed the highest larvicidal activity, and CL_50-90_ values were calculated. C2F7 was selected to carry out Column 3 and isolate a bioactive molecule. Column 3 had the same chromatographic conditions as Columns 1 and 2. Column 3 was separated into 26 samples; samples 13-15 crystallized into needle crystals. These crystals were decanted, and the resulting crystals were washed with *n-*hexane, yielding 256 mg.

### TLC and HPLC Analysis

Analytical TLC was carried out on a precoated Merck® silica gel 60F254 or RP-18F254 plates. Ceric sulphate reagent was used to visualize terpenes.

HPLC separations were performed on a Waters 2695 separations module equipped with a Waters 2996 photodiode array detector and HPLC analysis was carried out using a LiChrospher® 100 RP-18 column (4 mm × 250 mm, 5 μm) (Merck, Kenilworth, NJ, USA). The mobile phase consisted of two solvent reservoirs, A (H_2_O-Trifluoroacetic acid 0.05%) and B (CH_3_CN). The gradient system was as follows: 0–8 min, 100–0% B; 9–12 min, 90–10% B; 13–15 min, 80–20% B; 16–20 min, 70–30%, 21–25 min, 0–100% B, and 26–28 min 100–0% B. The flow rate was set at 1 mL/min, with a sample concentration of 2 mg/mL and an injection volume of 10 μL[19]. The absorption was measured at λ= 205 nm to visualize terpenes.

### CG-MS Analysis

The GC-MS analysis was performed using an Agilent Technologies HP 6890 gas chromatograph coupled to a quadrupole mass detector MSD 5973 (HP Agilent) and an HP-5MS capillary column (length: 30 m; inner diameter: 0.25 mm; film thickness: 0.25 μM). A constant flow of helium was set as the carrier gas to the column at 1 mL/min. The inlet temperature was fixed at 250 °C, while the oven temperature was initially kept at 40 °C for 1 min and increased to 280 °C at intervals of 10 °C/min. The mass spectrometer was used in positive electron impact mode with an ionization energy of 70 eV. Detection was performed in selective ion monitoring mode. The signals were identified and quantified using target ions. The compounds were identified by comparing their mass spectra with the NIST library version 1.7a. The relative percentages were determined by integrating the signals using GC Chem Station software, version C.00.01. The composition was reported as a percentage of the total signal area.

### NMR Experiments

One and two-dimensional Nuclear Magnetic Resonance (NMR) experiments (^1^H, COSY, HSQC, HMBC and DEPTq) were performed on a Bruker AVANCE III HD at 500 MHz. CD_3_COCD_3_ were used as solvents with tetramethylsilane (TMS) as an internal standard. Chemical shifts (δ) are reported in ppm values and coupling constants are in Hz.

### Near Infrared spectroscopy Analisys

NIR spectra were recorded on a Foss NIRSystems-6500 near infrared spectrophotometer (Raamsdonksveer, The Netherlands) following the methodology reported by López-Arellano et al. [20]

### Melting Point Experiment

A Fisher-Johns melting point apparatus was used to determine melting point. A small amount (less than 1 mg) of cristal was well spread placed between two coverslips. The heating control was setted at full power until 20 degrees of the theoretical melting point. Then, the power was setted to increase 1 °C per minute. The determination of the melting point was perfomed in triplicate.

### Statistical Analyses

Differences among lethality percentage means were compared using the Duncan test (*p* < 0.05). Lethal concentrations (LC_50_ and LC_90_) were determined with the PROBIT procedure included within the SAS statistic package.

## Results

### Yield extraction

A maceration extraction was performed using different solvents (methanol, ethyl acetate and *n-*hexane). The methanol extract (ME) showed a yield percentage of *4*.*10%*, the ethyl acetate extract (EAE) 3.86%, and the *n-*hexane extract (HE) 1.09%. ME exhibited the highest yield, followed by EA and HE, respectively.

### Anthelmintic activity of *Artemisia cina* crude extracts

The *in vitro* lethal concentration of the crude extracts against *Haemonchus contortus* infective larvae (L_3_) is shown in Table 1.

**Table 1.**
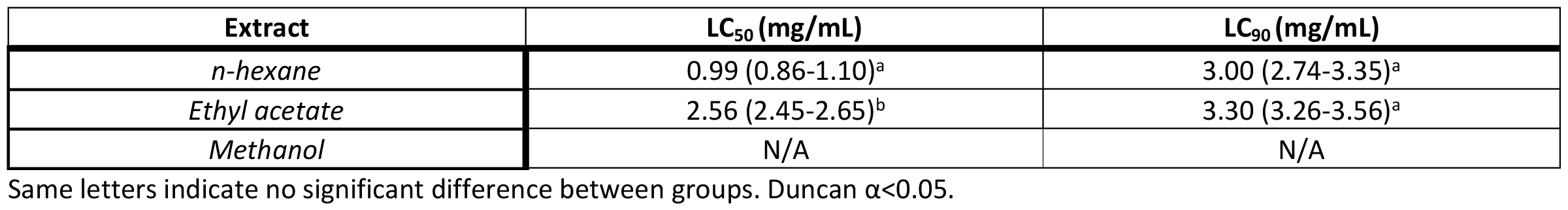
Lethal concentration of *Artemisia cina* extracts on *Haemonchus contortus* infective larvae L_3_.

| Extract | $\text{LC}_{50}$ (mg/mL) | $\text{LC}_{90}$ (mg/mL) |
| --- | --- | --- |
| *n*-hexane | 0.99 (0.86-1.10)<sup>a</sup> | 3.00 (2.74-3.35)<sup>a</sup> |
| Ethyl acetate | 2.56 (2.45-2.65)<sup>b</sup> | 3.30 (3.26-3.56)<sup>a</sup> |
| Methanol | N/A | N/A |

Same letters indicate no significant difference between groups. Duncan $\alpha < 0.05$ .

The HE had the best LC_50_, but no significative difference was found in LC_90_ compared with EAE. ME do not present a dose-response so, LC_50_ and LC_90_ cannot be calculated. EAE extract was chosen to perform the bio-guided separation due to the LC_90_ and the highest yield percentage of extraction, which is a crucial factor to consider in the formulation of pharmaceutic preparations.

### Bio-guided separation of the EAE of *A*. *cina* monitoring larvicidal activity against *Haemonchus contortus* infective larvae L_3_

EAE was separated into 61 samples and grouped into three fractions (C1F1, C2F2 and C3F3) based on their chemical similarity. C1F1, C2F2 and C3F3 were evaluated against L_3_ *H. contortus* infective larvae. C1F1 exhibited the highest larvicidal activity (Table 2), followed by C1F2. C1F3 did not exhibit at least 50% larvicidal activity; consequently, LC_50_ and LC_90_ were not calculated. Fraction C1F2 was chosen to continue with the separation process due to its significantly higher yield percentage compared to C1F1. The separation of C1F2 were grouped into 9 fractions, C2F7 displaying the highest larvicidal activity. C2F7 was then selected for attempts to isolate the bioactive molecule.

**Table 2.** *In-vitro* mortality percentage of *A. cina* fractions from bio-guided isolation of an anthelminthic compound against L_3_ *H. contortus* infective larvae.

| Treatments | Concentration (mg/mL) | Mortality % | LC<sub>50</sub>/LC<sub>90</sub> |
| --- | --- | --- | --- |
| Ethyl acetate extract | 8 | 100 ± 0 | 2.56 (2.45-2.65)<sup>c</sup> / 3.30 (3.26-3.66)<sup>c</sup> |
|  | 4 | 68.19 ± 11.86 |  |
|  | 2 | 21.85 ± 4.12 |  |
|  | 1 | 20.56 ± 12.10 |  |
|  | 0.5 | 21.90 ± 4.73 |  |
| C1F1 | 8 | 96.82 (± 2.39) | 0.72 (0.63-0.80)<sup>b</sup> / 2.58 (2.29-2.96)<sup>b</sup> |
|  | 4 | 93.83 (± 2.93) |  |
|  | 2 | 92.74 (± 3.24) |  |
|  | 1 | 90.74 (± 2.26) |  |
|  | 0.5 | 24.93 (± 5.17) |  |
|  | 0.25 | 32.73 (± 11.55) |  |
|  | 0.12 | 24.05 (± 3.58) |  |
|  | 0.62 | 28.24 (± 6.99) |  |
|  | 0.03 | 9.44 (± 3.30) |  |
| C1F2 | 8 | 95.28 (± 0.40) | 0.75 (0.60—0.89)<sup>b</sup> / 5.38 (4.69-6.38)<sup>d</sup> |
|  | 4 | 85.54 (± 2.92) |  |
|  | 2 | 78.78 (± 1.23) |  |
|  | 1 | 78.11 (± 7.38) |  |
|  | 0.5 | 27.85 (± 2.54) |  |
|  | 0.25 | 29.17 (± 7.55) |  |
|  | 0.12 | 30.34 (± 4.64) |  |
|  | 0.62 | 23.11 (± 10.71) |  |
|  | 0.03 | 26.55 (± 5.20) |  |
| C1F3 | 8 | 37.04 (± 7.07) | - |
|  | 4 | 21.24 (± 7.39) |  |
|  | 2 | 14.97 (± 2.11) |  |
|  | 1 | 14.63 (± 2.11) |  |
|  | 0.5 | 15.52 (± 2.09) |  |
| C2F1 | - | - |  |
| C2F2 | - | - |  |
| C2F3 | - | - |  |
| C2R4 | - | - |  |
| C2F5 | 0.75<br>5.38 | 83.78 ($\pm 5.17$)<br>82.41 ($\pm 2.57$) | |
| C2F6 | 0.75<br>5.38 | 61.87 ($\pm 6.81$)<br>77.02 ($\pm 4.71$) | |
| C2F7 | 0.75<br>5.38 | 80.84 ($\pm 3.99$)<br>92.58 ($\pm 0.90$) | |
| C2F8 | 0.75<br>5.38 | 60.59 ($\pm 9.44$)<br>70.91 ($\pm 1.69$) | |
| C2F9 | 0.75<br>5.38 | 18.39 ($\pm 2.35$)<br>21.26 ($\pm 3.36$) | |
| C2F7 | 6<br>3<br>1.5<br>0.75<br>0.37<br>0.19<br>0.93<br>0.04<br>0.02<br>0.01 | 91.21 ($\pm 1.55$)<br>87.80 ($\pm 2.36$)<br>83.61 ($\pm 3.30$)<br>76.08 ($\pm 5.16$)<br>69.21 ($\pm 1.37$)<br>65.00 ($\pm 9.40$)<br>70.66 ($\pm 5.54$)<br>44.25 ($\pm 2.60$)<br>30.17 ($\pm 4.65$)<br>21.11 ($\pm 3.08$) | 0.11 (0.13-0.90)<sup>b</sup> / 4.64 (3.64-6.08)<sup>c,d</sup> |
| C313-15P | 2<br>1<br>0.5<br>0.25<br>0.12<br>0.06<br>0.03<br>0.015 | 100 ($\pm 0.00$)<br>98.32 ($\pm 2.01$)<br>92.21 ($\pm 5.83$)<br>88.59 ($\pm 5.51$)<br>95.50 ($\pm 3.89$)<br>100 ($\pm 0.00$)<br>90.24 ($\pm 16.89$)<br>12.52 ($\pm 2.06$) | 0.01 (0.01-0.02)<sup>a</sup> / 0.21 (0.19-0.25)<sup>a</sup> |

Same letters indicate no significant difference between groups. Duncan $\alpha < 0.05$ .

In Column 3, where C2F7 was separated, a colorless needle crystal in samples 13-15 were observed. The larvicidal activity of compound 1 was determined, CL_50_ was 0.13 (0.11 - 0.14) mg/mL and CL_90_ 0.40 (0.37 - 0.44) mg/mL. In comparison with EAE larvicidal activity (CL_50_ 2.56 (2.45-2.65) mg/mL and CL_90_ 3.30 (3.26-3.66) mg/mL), compound 1 exhibited considerably more lethal activity against infective larvae.

In Fig. 1, is the schematic representation that illustrates the bio-guided separation methodology applied to the EAE, monitoring the larvicidal activity against the L_3_ of *H. contortus* infective larvae *in vitro*. As a result, an unknown compound with anthelmintic activity was isolated, yielding 0.01% in comparison to the EAE.

**Figure 1.**
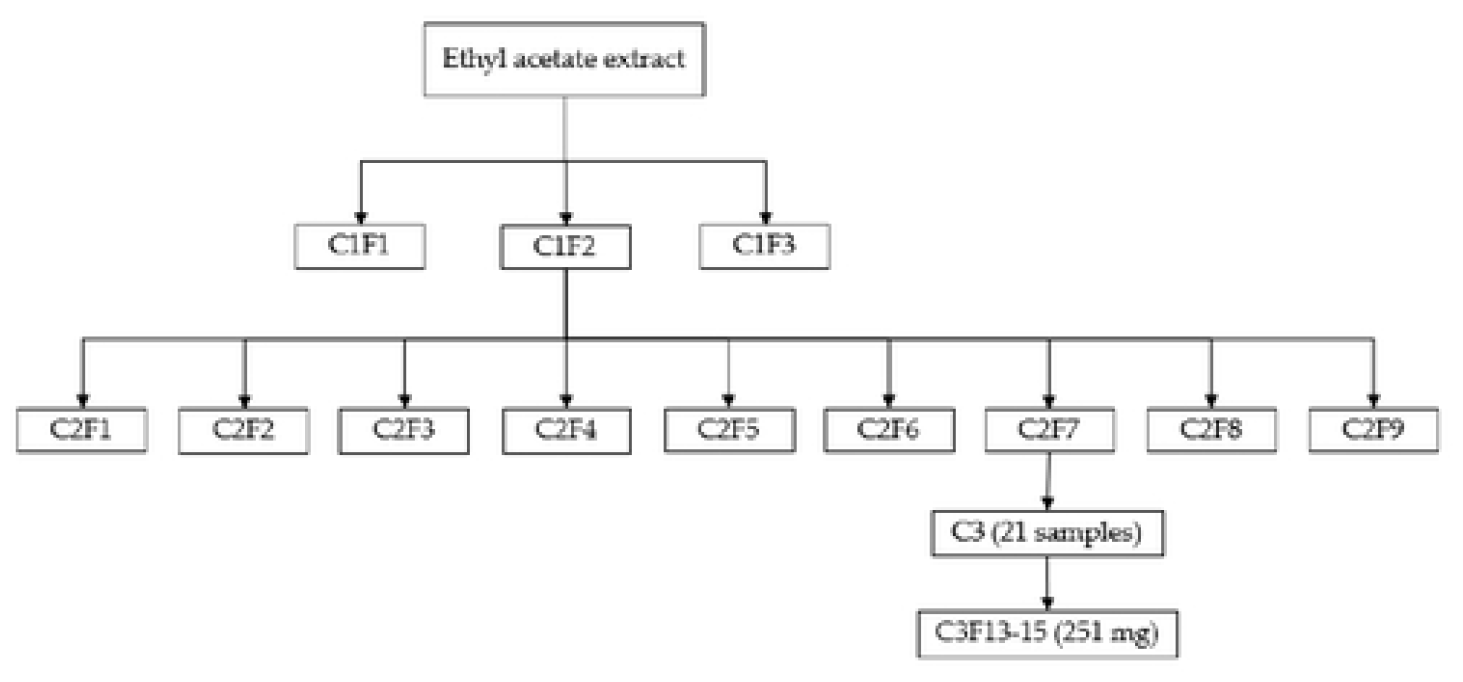
Bio-guided separation scheme of the ethyl acetate crude extract of *Artemisia cina* monitoring larvicidal activity on L_3_ *Haemonchus contortus* infective larvae.

### Identification of compound 1

Compound 1 was obtained through the chromatographic separation of C2F7, using 1:1 *n-*hexane-ethyl acetate as mobile phase. Colorless needle crystals were obtained and according to UV and mass spectra, there was no information of the compound reported in *Artemisia cina*. It was necessary to perform one- a two-dimensional NMR spectroscopy to identify it.

This compound was soluble in dichloromethane-metanol 1:1. TLC showed a weak florescent band when observed under λ = 254 nm UV light and no fluorescence at λ = 365 nm. HPLC analysis showed a peak at 15.775 min and an absorption spectrum λ = 211.0 nm, typically of terpenes (Fig. 2), and a [M+H]^+^= 283 m/z (S1 Fig.)The NIRS spectra (S2 Fig.) exhibited 2060 nm R-OH combination, 2116 nm C-C combination. The presence 1118 nm CH_3_ third overtone region, 1406 CH_3_ and CH_2_ second overtone region, 1690 nm, 1714 nm and 1738 nm CH_3_, CH_2_ and CH first overtone region. 2270 nm, 2290 nm, 2310 nm CH_3_, CH_2_ and CH combination area [9]. Also displayed 213 °C melting point.

**Figure 2.**
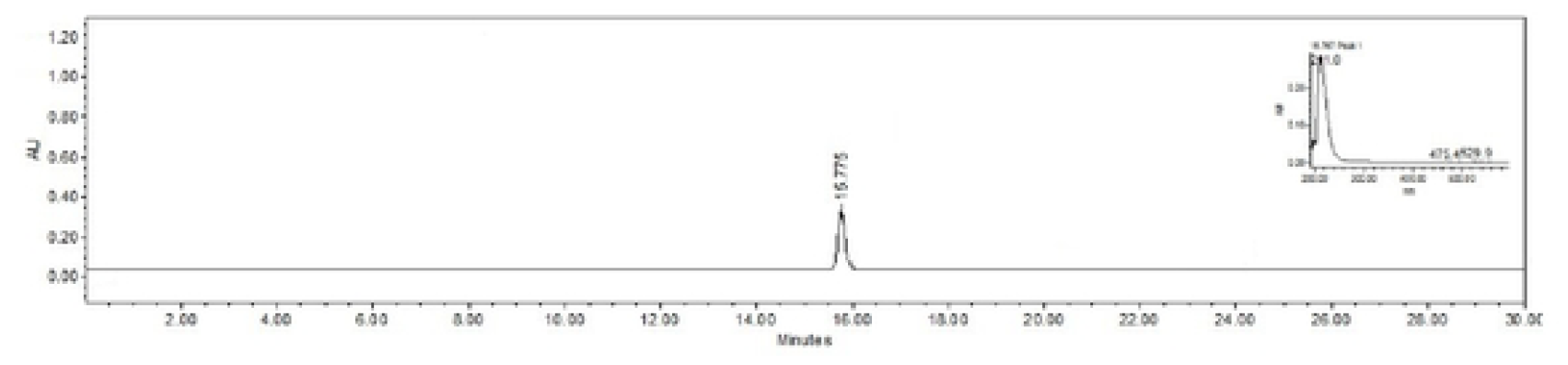
Chromatogram and UV-VIS spectra of C3 13-15P, obtained by HPLC-DAD and viewed at 215 nm.

**Figure 3.**
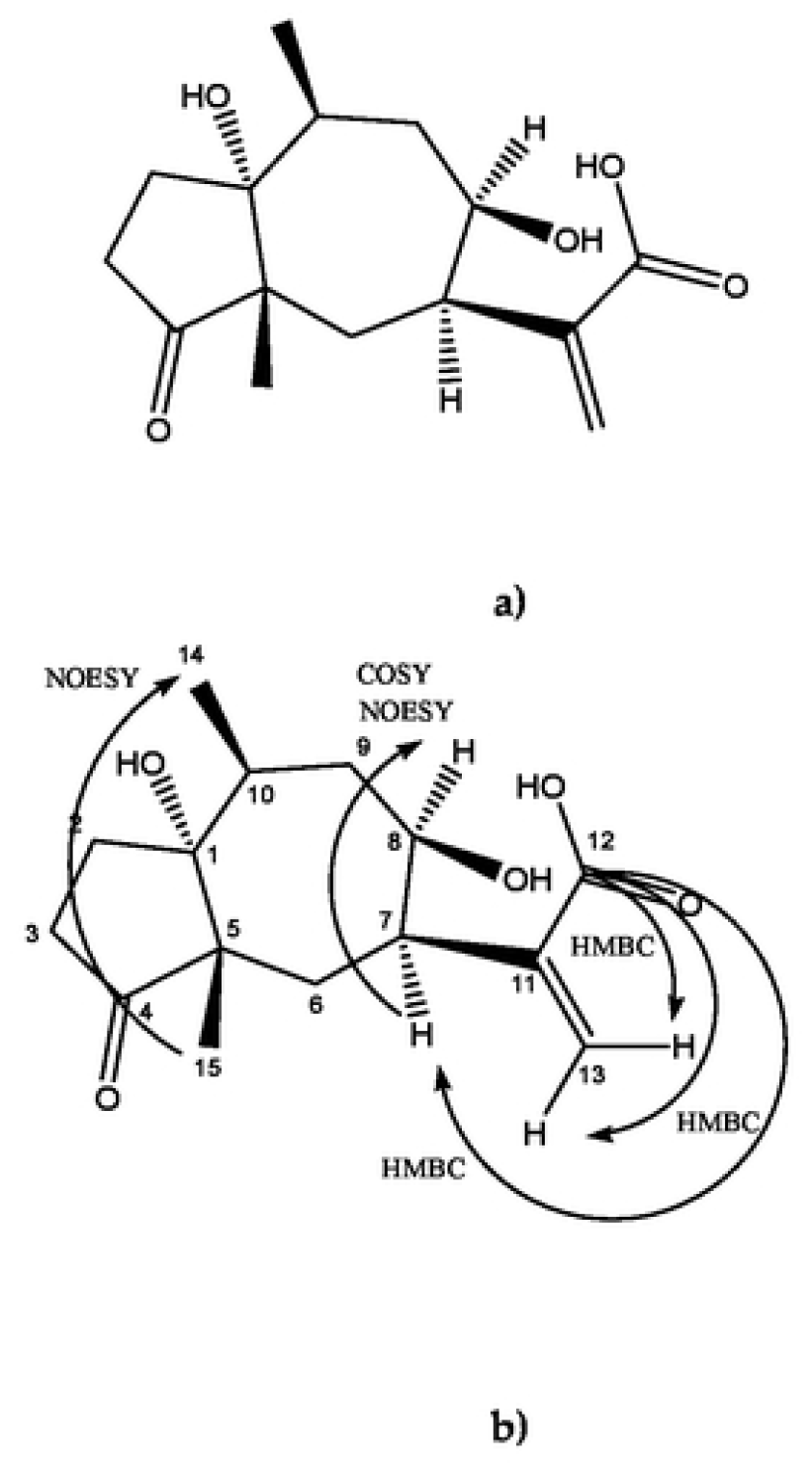
**New sesquiterpene isolated from aerial parts of** *Artemisia cina* **(a) Structure of cinic acid (b) Correlations of cinic acid (HMBC, COSY and NOESY at 500MHz)**

^1^H-NMR spectrum (Table 3 and S3 Fig.), indicated de presence of two methyl groups (δ 1.20 d, J= 6.83 Hz) and (δ 1.06 s), one CH-O proton (δ 4.27 m, br) and Ha (δ 6.21 t, J= 1.01, 1.01 Hz), Hb (δ 5.72 dd, J= 0.91, 1.70 Hz) of a terminal alkene. The ^13^C - NMR spectrum (Table 3) showed 15 signals, typically of sesquiterpenes. This was correlated with the absorption spectrum. The δ 216.56 signal (C-4) correspond to a ketone carbonyl and δ 168.20 to an ester or carboxylic acid carbonyl. The δ 90.38 and 74.84 corresponds to C-O signals. The δ 141.36 (C-11) and δ 125.07 (C-13) signals were alkene type, aromatic signal was discarded due to de absence of signals between δ 7.00 and δ 8.00 in the ^1^H -NMR spectrum. Methyl groups were observed at δ 21.15 (C-14) and δ 15.06 (C-15).

**Table 3.** ^1^H -NMR and ^13^C -NMR spectroscopy data of compound 1 (CD_3_COCD, 500 MHz)

| Position | DEPTq | $^1H$ (J in Hz) | $^{13}C$ |
| --- | --- | --- | --- |
| 1 | C | - | 90.38 |
| 2 | $CH_2$ | $\alpha$ 1.98 (1H, dd, J= 13.8, 5.2)<br>$\beta$ 1.63 (1H, br) | 28.43 |
| 3 | $CH_2$ | $\alpha$ 2.37 (1H, m)<br>$\beta$ 2.37 (1H, m) | 27.42 |
| 4 | C | - | 216.56 |
| 5 | C | - | 53.60 |
| 6 | $CH_2$ | 2.44 (1H, dd, J = 9.8, 1.4)<br>2.29 (1H, m) | 34.11 |
| 7 | CH | 2.85 (1H, m, br) | 39.96 |
| 8 | CH | 4.27 (1H, m, br) | 74.84 |
| 9 | $CH_2$ | 2.14 (1H, m)<br>1.42 (1H, m) | 34.44 |
| 10 | CH | 2.14 (1H, m) | 41.51 |
| 11 | C | - | 141.36 |
| 12 | C | - | 168.20 |
| 13 | $CH_2$ | Ha 6.21 (1H, t, J= 1.01, 1.01),<br>Hb 5.72 (1H,dd, J= 0.91, 1.70) | 125.07 |
| 14 | $CH_3$ | 1.24 (3H, d, J=7.00) | 15.06 |
| 15 | $CH_3$ | 1.06 (3H, s) | 21.15 |

Sesquiterpene lactones are commonly found in *Artemisia* genus [10]. The presence of ketone (δ 216. 56), ester carbonyl (δ 168.20) and alkene (δ 141. 36 and δ 125.07) signals are frequently of guaianolides and pseudoguaianolides (5-7 bicyclic compounds) [11]. The presence of pseudoguaianolide could be confirmed due to the presence with a methyl group at the C-5 ring junction (C-14 δ 21.15) and C-10 (C-15 δ 15.06). The signals δ 34.11 and δ 34.44 signals, correspond to CH_2_ with similar electronic environment corresponding to C-6 and C9 sesquiterpene lactone lactonized towards C7-C8. Surprisingly, HMBC analysis did not show correlation between C-12 and H-8 (δ 4.27 m, br) but had correlation with H13a (δ 6.21 t, J= 1.01, 1.01 Hz) H13b (δ 5.72 dd, J= 0.91, 1.70 Hz)) and H7 (δ 2.85 m, br) (S5 Fig.), so it was not lactonized and the ring is open. COSY and NOESY also show the correlation between H7 (δ 2.85 m, br) and H8 (δ 4.27 m, br) consistent with *cis*-orientation of the protons (S6 Fig and S7a Fig.). According to the NOESY, the two methyl groups (δ 1.20 d, J= 6.83 Hz) and (δ 1.06 s)) are *cis*-orientated too (S7b Fig.). The proposed structure is shown at Fig.

Compound 1 was identified 11-[(1*R*,5*S*,7*R*,8*R*,10*S*,)-1,8-dihydroxy-5,10-dimethyl-4-oxodecahydroazulen-7-yl] acrylic acid, not previously reported and was named cinic acid. Due to the structural similarity to the pseudoguainolides, the compound was numbered accordingly.

## Discussion

*Haemonchus contortus* is a hematophagous gastrointestinal parasite that posses a threat to grazing sheep. Its feeding habits contribute to conditions such as anemia and poor digestion, potentially leading to mortality in young individuals. In adults, chronic inflammation, weight loss, and persistent diarrhea are common, resulting in significant losses in animal production worldwide [12]. Furthermore, the presence of drug-resistant helminths poses a significant challenge to the sustainability of current helminth control strategies [13], It is imperative to develop alternative antiparasitic treatments against *H. contortus*, including the exploration of plant-based anthelmintics through discovery and development efforts.

*Artemisia* species have attracted significant research attention due to the presence of sesquiterpenoid lactones, coumarins, flavonoids, and phenolic acids. These compounds are responsible for a wide range of biological activities, including hepatoprotective, neuroprotective, antidepressant, cytotoxic, antitumor, digestion-stimulating and antiparasitic effects [10]. The *n-*hexane (*n*-HE) extract of *Artemisia cina* has been documented for its anthelmintic activity against *H. contortus*, targeting eggs and L_3_ infective larvae [8], transitional larvae L_3_-L_4_ [4], and naturally infected periparturient goats [3]. Those activities are attributed to the presence of two lignans, 3′-Demethoxy-6-O-Demethylisoguaiacin and norisoguaiacin [8]. However, the low yield percentage of the *n*-HE and the lignans make it challenging to use for formulating or creating *A. cina*-based pharmaceutical preparations. In this study, the ethyl acetate extract (EAE) was chosen for its significantly higher extraction yield (3.86 times higher) compared to the *n*-HE, while maintaining a similar larvicidal activity to *n*-HE.

The bio-guided isolation of the EAE from *A. cina*, led to separate and identify a new sesquiterpene that was called cinic acid. This compound exhibits a great larvicidal activity, was 256 times more active at LC_50_ and 15.71 times at LC_90_ compared to EAE, likely responsible for a significant portion of the overall extract activity. The compound yields 0.01% relative to the extract but is possible the presence of synergism between cinic acid and other compounds of the EAE but this hypothesis had to be probe.

Sesquiterpenes are secondary metabolites with a 15-carbon skeleton built from three isoprene units. They are commonly cyclized and found in the Asteraceae family and exhibits several pharmacological activities [14]. The predominant sesquiterpenes isolated in *Artemisia* species are sesquiterpene lactones [15]. Sesquiterpene lactones are categorized into four primary groups: germacranolides (with a 10-membered ring), eudesmanolides (6–6 bicyclic compounds), guaianolides, and pseudoguaianolides (5–7 bicyclic compounds). A distinguishing characteristic of sesquiterpene lactones (STLs) is the existence of a γ-lactone ring closed either at C-6 or C-8. This γ-lactone often includes, in numerous instances, an exo-methylene group that is conjugated to the carbonyl group [16]. Cinic acid is a novel sesquiterpene with a structure resembling that of a pseudoguaianolide-type sesquiterpene lactone, specifically an ambrosanolide. It features two methyl groups *cis*-orientated at C-5 and C-10 and an exocyclic methylene conjugated to a γ-carbonyl moiety necessary for the biological activity. The notable distinction lies in the absence of lactonization at C8. The activity of sesquiterpene lactones is determined by the presence of the α-methylene, γ-lactone system, which acts as a Michael acceptor, allowing interaction with thiol groups of proteins [17]. The α-methylene, γ-carbonyl system present in cinic acid could explain its *in vitro* anthelmintic activity against L_3_ infective larvae of *H. contortus* [18].

## Conclusion

The bio-guided separation of ethyl acetate extract (EAE) allowed the identification of a new sesquiterpene with anthelmintic activity against *H. contortus* L_3_ infective larvae. The EAE could be a promising candidate for a plant-based pharmaceutical preparation with anthelmintic activity from *Artemisia cina*.

## Supporting information

**S1 Fig**. Mass spectra of cinic acid obtained by CG-MS.

**S2 Fig**. Cinic acid near infrared spectroscopy spectra.

**S3 Fig**. ^1^H NMR spectra of cinic dissolved in CD_3_COCD_3_ and obtained at 500 MHz.

**S4 Fig**. DEPTq spectra of cinic acid dissolved in CD_3_COCD_3_ and obtained at 500 MHz.

**S5 Fig**. HMBC spectra of cinic acid dissolved in CD_3_COCD_3_ and obtained at 500 MHz.

**S6 Fig**. COSY experiment of cinic acid dissolved in CD_3_COCD_3_ and obtained at 500 MHz a) COSY spectra of cinic acid and b) COSY spectra of the correlation between H7 (δ 2.85 m, br) and H8 (δ 4.27 m, br) consistent with *cis*-orientation of the protons.

**S7 Fig**. NOESY experiment of cinic acid dissolved in CD_3_COCD_3_ and obtained at 500 MHz a) NOESY spectra of cinic acid and b) NOESY spectra of the correlation between the two methyl groups (δ 1.20 d

## Acknowledgments

This research was funded by the Support Program for Research and Technological Innovation Projects (PAPIIT-UNAM), titled: Evaluación del efecto tóxico del extracto *n*-hexánico de *Artemisia cina* y cinaguaiacina sobre los parámetros bioquímicos en sangre y alteraciones anatomopatológicas en ratas Wistar después de su administración por vía oral (IA204822). Some materials were covered with resources from CIBIS-IMSS. LDAP was supported by Postdoctoral Grants Program from National Autonomous University of Mexico. To the social service students who supported in obtaining biological material: Eduardo Rico-Mejía, Fernanda Lazcano Cárdenas and Lian Guillermo Cortés Campos.

**Acknowledgments**

**Conceptualization:** Higuera-Piedrahita Rosa Isabel, López-Arellano Raquel, Arango de la Pava Luis David

**Data curation:** Manasés González-Cortázar, Alejandro Zamilpa

**Formal analysis:** Cuéllar-Ordaz Jorge Alfredo, González-Cortázar Manasés, Zamilpa Alejandro, De la Cruz-Cruz Héctor Alejandro

**Funding acquisition:** Higuera-Piedrahita Rosa Isabel, López Arellano Raquel

**Investigation:** Higuera-Piedrahita Rosa Isabel, López-Arellano Raquel, Arango de la Pava Luis David

**Supervision:** Higuera-Piedrahita Rosa Isabel, González-Cortázar Manasés, Zamilpa Alejandro

**Validation:** Manasés González-Cortázar, Alejandro Zamilpa

**Writing – original draft:** Arango de la Pava Luis David, Manasés González-Cortázar, Alejandro Zamilpa, De la Cruz-Cruz Héctor Alejandro

**Writing – review & editing:** Arango de la Pava Luis David, Manasés González-Cortázar, Alejandro Zamilpa, De la Cruz-Cruz Héctor Alejandro, Cuéllar-Ordaz Jorge Alfredo, López-Arellano Raquel, Higuera-Piedrahita Rosa Isabel

## Notes

### Competing Interest Statement

The authors have declared no competing interest.

